# Mendelian randomizations with spatial gene networks reveal shared and distinct drivers of risk in major skin cancer types

**DOI:** 10.1101/2025.02.25.640031

**Authors:** Michael Pudjihartono, Justin M. O’Sullivan, William Schierding

**Author notes:** Corresponding author: William Schierding; and Justin O’Sullivan.

## Abstract

Skin cancer is the most common malignancy worldwide, comprising three major types: melanoma, basal cell carcinoma (BCC), and squamous cell carcinoma (SCC). A critical goal in advancing our understanding of skin cancer is to identify the shared and distinct biological mechanisms that drive the risk profiles for each skin cancer type. In this study, we integrated tissue-specific chromatin conformation and expression quantitative trait loci data (spatial eQTL) to construct gene regulatory networks (GRNs) for melanocytes, sun-exposed skin, not sun-exposed skin, and blood. These GRNs, which capture spatially regulated gene expressions, were used as instrumental variables along with GWAS summary statistics for melanoma, BCC and SCC, to infer causal relationships between specific gene expression changes and each type of skin cancer. These Mendelian randomization analyses identified 82, 62, and 125 causal genes for melanoma, BCC, and SCC, respectively, with many of these genes not evident from the GWAS data alone. Our analyses revealed distinct mechanisms for each skin cancer type: telomere maintenance and nevus pathways in melanoma, inherited immune traits in BCC, and p53 dysfunction in SCC. Notably, apoptosis and pigmentation emerged as shared biological processes across skin cancers. These findings provide new insights into the genetic drivers of three distinct skin cancers, highlighting new gene targets which can be used to increase diagnostic precision, as well as potential therapeutic targets.

## 1 Background

Skin cancer is the most prevalent human malignancy^1^. The three major types—melanoma, basal cell carcinoma (BCC), and squamous cell carcinoma (SCC)—are experiencing increasing worldwide incidence^1–3^. Among these, BCC and SCC are the most common, incurring substantial quality of life and financial burden—ranking among the top five most expensive malignancies to treat in the USA^4^. However, melanoma accounts for the majority of skin cancer-related deaths^5,6^ and has recently become the most rapidly increasing cancer in white populations^7^. Identifying the shared and distinct biological mechanisms that define the etiology of each skin cancer type, including those contributing to increased inherited risk^8^, is critical to advance our understanding of these diseases.

Genome-wide association studies (GWAS) have identified dozens of genomic loci associated with the risk of developing each type of skin cancer^9–11^. These GWAS provide a framework for identifying putative risk loci, but due to the nature of these association studies, they are not aimed at identifying causal genes. This is due to several reasons: 1) the vast majority of single nucleotide polymorphisms (SNPs) identified by GWAS are located in the non-coding regions of the genome^12^; 2) the complicated linkage disequilibrium (LD) structure of the genome means that the tag GWAS SNP is not necessarily causal^13^; and 3) SNPs can affect phenotype via distant regulation of gene expression, which limits the effectiveness of conventional nearest-gene approaches for identifying SNP functional targets^14^. Identifying the functional targets of melanoma, BCC, and SCC-associated loci—including both shared and unique—will enable the discovery of genes and pathways involved in the aetiology of each skin cancer type. This, in turn, will aid in the development of personalized therapies, drug repurposing, and prevention strategies.

We previously investigated how melanoma-associated SNPs influence gene expression by integrating tissue-specific chromatin interaction (Hi-C) and expression quantitative trait loci (eQTL) data^15^. While these findings provided valuable insights, they have yet to establish statistical evidence supporting the causal involvement of these genes in melanoma pathogenesis. Mendelian randomization (MR) is a statistical method that uses genetic variants as instrumental variables to estimate the causal effect of a modifiable exposure on a disease outcome^16^. When appropriately designed, MR is less susceptible to common epidemiological biases and can yield reliable causal inferences^17^.

In this study, we first constructed gene regulatory networks (GRNs) by integrating tissue-specific Hi-C and eQTL data^15,18^. These GRNs consist of SNPs that physically interact with and regulate the expression of specific target genes, referred to as “spatial eQTLs”. We then performed two-sample MR^19^, using these spatial eQTLs as instrumental variables, along with GWAS summary statistics for melanoma, BCC and SCC, to infer causal relationships between specific gene expression changes and each type of skin cancer. These analyses identified 82, 62, and 125 causal genes for melanoma, BCC, and SCC, respectively. Pathway analyses revealed a unique enrichment of telomere maintenance and nevus-related genes in melanoma, while BCC showed a prominent enrichment of immune genes influencing its susceptibility. The findings in SCC were consistent with p53 dysfunction in its early pathogenesis. Overall, our study provides new insights into the shared and distinct biological mechanisms driving skin cancer development.

## 2 Materials and methods

### 2.1 Construction of tissue-specific gene regulatory networks

Gene regulatory networks (GRNs) were constructed by analyzing all common SNPs (MAF ≥ 0.05; n=∼40×10^6^) for spatial eQTL associations using the CoDeS3D algorithm^20,21^. Briefly, Hi-C chromatin contact data were utilized to identify distal DNA fragments that are spatially connected to each SNP. If a distal fragment overlapped a protein-coding gene region (as per the GRCh38 gene annotations from GENCODE v26^22^) in at least two replicates of the Hi-C data, a spatial SNP-gene pair was established. For each pair, eQTL data were then used to determine whether the SNP was also associated with expression level changes of the target protein-coding gene, thereby establishing a spatial eQTL-gene association. Spatial eQTL-gene associations were considered significant if they passed a significance threshold of *P* ≤ 1×10^-5^. Tissue-specific GRNs were generated for four tissues: melanocyte, sun-exposed skin, not sun-exposed skin, and blood using Hi-C data from three skin and four blood cell lines, along with eQTL data from three GTEx^23^ and one melanocyte^24^ dataset (Supplementary Table 1).

### 2.2 Mendelian randomization analyses

To identify potentially causal genes for each type of skin cancer, we conducted two-sample mendelian randomization (MR) analyses using the TwoSampleMR R package (v0.5.6)^25^ (see STROBE-MR Supplementary Checklist^26^). MR is a statistical method that uses genetic variants (*i.e.*, SNPs) as “instrumental variables” to estimate the causal effect of an exposure (*i.e.*, gene expression) on an outcome (*i.e.*, different type of skin cancer). The instrumental variables used in MR must satisfy three assumptions: 1) the instrumental variables are associated with the exposure of interest; 2) the instrumental variables are independent of any potential confounders; and 3) the instrumental variables influence the outcome solely through their association with the exposure (*i.e.*, no horizontal pleiotropy)^16^.

Spatial eQTL-target gene pairs (*P* ≤ 1×10^-5^) from each GRN were used as the exposure dataset. Palindromic SNPs (*i.e*., those with alleles that are reverse complements, such as A/T or C/G) were removed to prevent distortion of strand orientation or allele coding during the harmonization process^27–30^. To ensure the independence of instrumental variables for each exposure, we performed LD clumping within a 10 Mb window and r^2^ cut-off of < 0.001 based on the 1000 Genomes project European population^31^. For melanoma, we used GWAS summary statistics for 36,760 cases and 375,188 controls from Landi et al.^11^ as the outcome data. For BCC and SCC, we used GWAS summary statistics from Adolphe et al.^9^ (17,416 cases, 375,455 controls) and Seviiri et al.^10^ (7,402 cases, 286,892 controls) as the outcome data, respectively. Each pairwise exposure and outcome data were then harmonized to ensure that the effect of a spatial eQTL on both outcome and exposure is relative to the same allele (Supplementary File 1). Once harmonized, MR was performed using the Wald ratio method for exposures with one instrumental variable, inverse variance weighted (IVW) method for exposures with two instrumental variables, and weighted median method for exposures with ≥3 instrumental variables^32^. Target genes with MR *P* value ≤ the Bonferroni-correction threshold of 0.05/(number of unique exposure genes) were considered statistically significant.

We performed several sensitivity analyses for target genes with multiple instrumental variables. First, Cochran’s Q test was performed; instrumental variables with *P* value < 0.05 were considered heterogeneous. For target genes with ≥3 instrumental variables, MR-Egger regression was further performed to detect horizontal pleiotropy. A significant deviation from zero in the intercept (*P* value < 0.05) was taken as evidence of horizontal pleiotropy. Genes failing any of these sensitivity tests were excluded from the list of significant causal genes.

### 2.3 Determination of independent loci

Across four tissues and three skin cancer types, all unique significant instrumental variable SNPs were then grouped into independent loci using PLINK^33^, with LD pruning performed based on the 1000 Genomes project European population. The pruning process involved sorting the SNPs by genomic coordinates and applying a sliding window of 50 SNPs, which shifted by 5 SNPs at each step. Within each window, SNP pairs exhibiting an r^2^ > 0.2 were identified, and one SNP from each pair was pruned out (removed) to ensure that the remaining SNPs were approximately independent. This procedure resulted in the identification of the set of independent (tag) SNPs, each tagging an independent locus. Each non-tag SNP in the original dataset was then assigned to the locus tagged by the tag SNP with which it had the highest LD, given this was above an r^2^ of 0.2. This process organized all 319 SNPs into 144 independent loci (tagged by 144 independent SNPs) (Supplementary Table 2).

### 2.4 Identification of genes previously associated with skin cancer

Genes previously associated with skin cancer were identified using DisGeNET^34^ and the GWAS Catalog^35^. We downloaded the gene-disease association summary from DisGeNET on 16^th^ July 2024, for melanoma (UMLS CUI: C0025202), BCC (UMLS CUI: C0007117), and SCC (UMLS CUI: C0007137). Gene-disease associations were considered only from the “CURATED” source in DisGeNET. Additionally, we downloaded all associations from the GWAS Catalog on 16^th^ July 2024, for melanoma (EFO ID: EFO_0000756), BCC (EFO ID: EFO_1000529), and SCC (EFO ID: EFO_1000073). Skin cancer-associated genes were identified from the “MAPPED_GENE” column.

### 2.5 Permutation test for overlapping causal gene sets

We conducted permutation tests to assess the significance of overlap among causal gene sets for melanoma, BCC, and SCC. The gene universe consisted of unique genes from the four tissue-specific GRNs (n = 11,833). For each skin cancer type, we generated 100,000 random samples, drawing gene sets equal in size to the observed number of causal genes identified in the MR analyses. In these simulations, each gene’s likelihood of being sampled was proportional to the number of GRNs (out of four) in which it was present. Specifically, for each gene, this likelihood was calculated by dividing the number of GRNs in which the gene was present by the total sum of tissue counts across all genes in the universe. This calculation normalized the probabilities to sum to 1, ensuring they could be interpreted as sampling probabilities. Genes were then sampled without replacement based on these normalized probabilities. Overlap was calculated for the combinations: melanoma and BCC, melanoma and SCC, BCC and SCC, and all three. *P* values were determined by the proportion of permutations where the overlaps were equal to or greater than those observed:

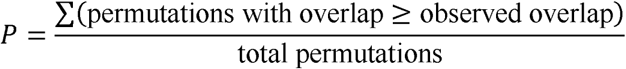

### 2.6 Functional enrichment analyses

Functional enrichment analysis was performed on each set of overlapping causal genes using the gprofiler2 R package^36^. The analysis evaluated genes for enrichment against biological terms derived from several ontologies and databases, including Gene Ontology (GO) biological processes (GO:BP), GO molecular functions (GO:MF), GO cellular components (GO:CC), Human Phenotype Ontology (HPO), and Reactome pathways (REAC). Significance of enrichment was determined using a Benjamini-Hochberg corrected *P* value threshold of < 0.05.

## 3 Results

### 3.1 Gene regulatory networks identify spatially constrained eQTLs in melanocyte, sun-exposed skin, not sun-exposed skin, and blood

Tissue-specific gene regulatory networks (GRNs) were constructed for melanocytes, sun-exposed skin, not sun-exposed skin, and blood by integrating chromatin interaction (Hi-C) data from three skin and four blood cell lines, along with eQTL data from three GTEx^23^ and one melanocyte^24^ dataset (Supplementary Table 1). The GRN of sun-exposed skin, not sun-exposed skin, and blood are comparable in size; however, the melanocyte GRN is notably smaller (Figure 1). This difference may reflect the relatively modest sample size of the melanocyte eQTL dataset (n = 106) compared to those for sun-exposed skin (n = 605), not sun-exposed skin (n = 517), and blood (n = 670). Overall, this approach allowed us to assign putative functions to SNPs by linking them to their target genes based on physical interactions and regulatory influence on gene expression (termed “spatial eQTLs”).

**Figure 1.**
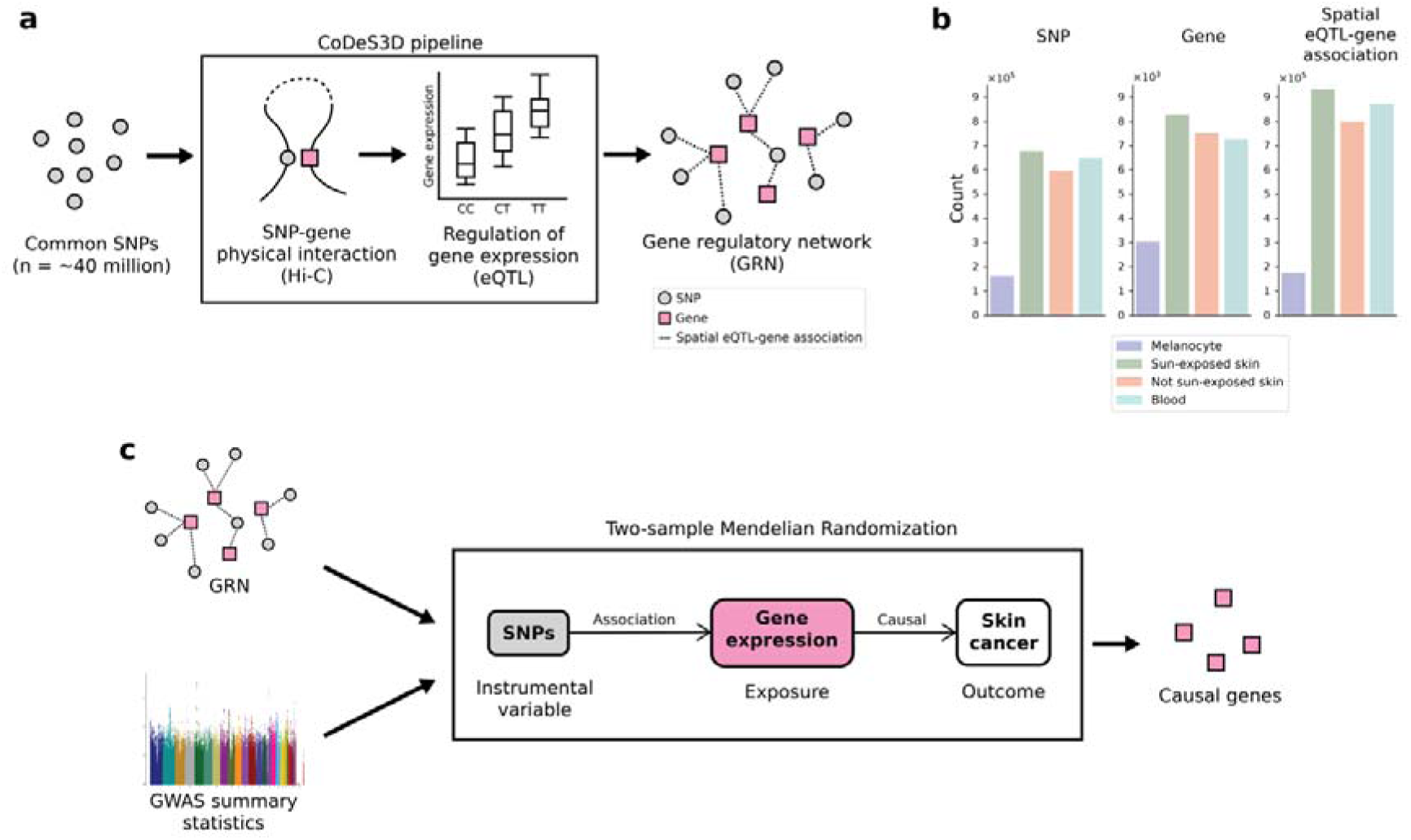
Overview of the analytical approach used in this study. (a) Gene regulatory networks (GRNs) were generated by integrating tissue-specific Hi-C chromatin contact data with eQTLs through the CoDeS3D algorithm, establishing spatial eQTL-gene associations for melanocyte, sun-exposed skin, non-sun-exposed skin, and blood tissues. (b) A comparison of the numbers of SNPs, genes, and spatial eQTL-gene associations identified in each tissue-specific GRN. (c) Two-sample Mendelian randomization analyses utilizing spatial eQTLs from each GRN as instrumental variables to infer causal relationships between gene expression and skin cancer outcomes.

### 3.2 Mendelian randomization identifies 82 genes causally associated with melanoma

We performed two-sample MR analyses to identify genes whose expression changes were putatively causal for melanoma (hereafter referred to as “causal genes”). We used spatial eQTLs from each GRN as instrumental variables and melanoma GWAS summary statistics^11^ as the outcome data (Figure 1c). Analyses were conducted according to the STROBE-MR guidelines^26^ (Supplementary Checklist). After removing genes that did not pass sensitivity tests (Supplementary Table 3), we identified 28 causal genes regulated by 25 spatial eQTLs within 22 independent loci in melanocyte, 42 causal genes in sun-exposed skin (41 spatial eQTLs; 28 loci), 44 causal genes in not sun-exposed skin (45 spatial eQTLs, 29 loci), and 36 causal genes in blood (36 spatial eQTLs; 29 loci) (Supplementary Table 4).

Unsupervised hierarchical bi-clustering of the MR β estimates highlighted patterns in the causal effects of genes across the four tissues (Figure 2a). The two skin tissues (sun exposed and not sun-exposed) exhibited the most similar patterns of causal gene effects, followed by melanocyte, with blood showing the most distinct patterns (Figure 2a). These differences align with the functional differences among these tissues, as melanocytes are a component of the skin, while blood represents a different tissue type. The unique pattern observed in blood may also be influenced by the use of distinct Hi-C cell lines compared to the other GRNs (Supplementary Table 1). Across the four tissues, we identified a total of 82 causal genes regulated by 128 spatial eQTLs in 57 independent loci (Figure 2b and c). Of these, 44 genes (54%) were shared between at least two tissues, including six genes (*CASP8*, *MTAP*, *PARP1*, *STN1*, *UQCC1*, and *MMP24-AS1*) that were shared across all tissues (Figure 2a). The causal effect directions were collinear for most of these shared genes (41/44; 93%) (Figure 2c).

**Figure 2.**
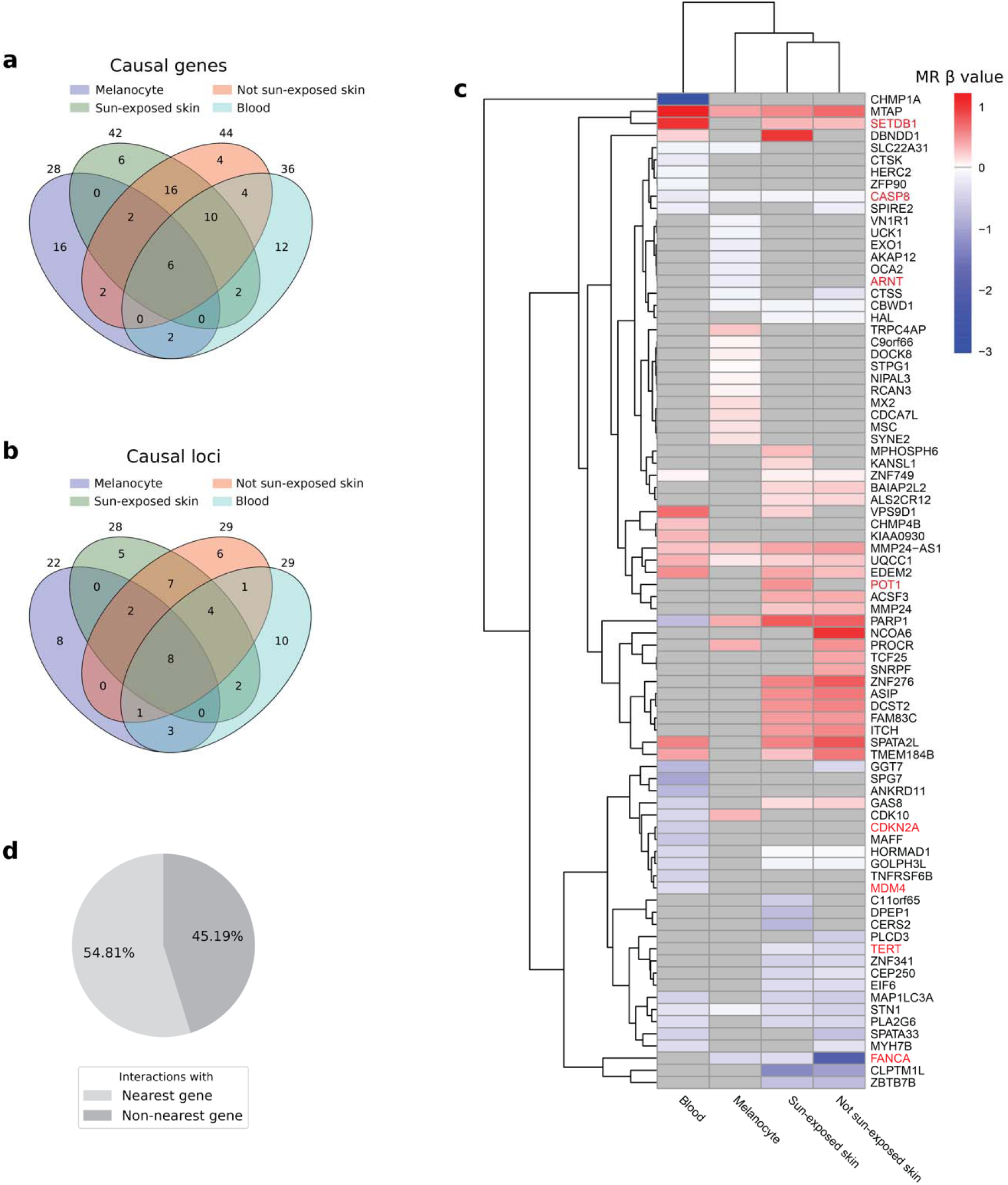
82 genes are causally associated with melanoma by 2-sample Mendelian randomization (MR). (a) Venn diagram showing the number of melanoma causal genes identified in each tissue. (b) Venn diagram showing the number of melanoma causal loci in each tissue. (c) Unsupervised hierarchical bi-clustering of the MR β estimates of causal genes across the four tissues. In the context of MR, the β value represents the effect of each unit increase in gene expression on the log(odds) of developing the disease. Therefore, a positive value (red) signifies that increased expression of a gene is associated with elevated melanoma risk, whereas a negative value (blue) denotes the opposite effect. Genes highlighted in red are recognized as known cancer drivers in the Cancer Gene Census database. (d) Pie chart showing the proportions of causal SNP-gene interactions that involve nearest genes versus non-nearest genes.

We identified a set of the causal genes (n = 38) that have not been previously associated with melanoma, either in DisGeNET^34^ or through the “nearest gene” approach represented in the GWAS Catalog^35^ (Supplementary Table 5). Additionally, 45% of the causal spatial eQTL-gene interactions involved non-nearest genes [Figure 2d]. Of note, among the causal genes, two melanoma driver genes, *CDKN2A* and *TERT*, were identified. This is a significant enrichment (hypergeometric *P* = 0.023) when compared to the driver genes (n = 56) listed in the Cancer Gene Census (CGC) database^37^ (Supplementary Table 6). Additionally, when considering all cancer driver genes (including those that have not been specifically annotated as a melanoma driver in the CGC; n = 737) six more genes (*CASP8*, *ARNT, FANCA*, *MDM4*, *POT1*, *SETDB*) were identified (hypergeometric *P* = 0.011; Supplementary Table 5, Figure 2a).

### 3.3 Mendelian randomization identifies 62 causal genes for BCC and 125 causal genes for SCC

We extended our two-sample MR analyses to include BCC and SCC, using their respective GWAS summary statistics^9,10^ as the outcome data. After removing genes that did not pass sensitivity tests (Supplementary Table 7), we identified a total of 62 causal genes regulated by 107 spatial eQTLs within 54 independent loci in BCC (Supplementary Figure 1a-d, Supplementary Table 8). For SCC, following sensitivity test exclusions (Supplementary Table 9), we identified 125 causal genes, regulated by 183 spatial eQTLs across 81 independent loci (Supplementary Figure 2a-d, Supplementary Table 10).

Unsupervised hierarchical bi-clustering of the MR β estimates across the four tissues revealed patterns similar to those seen in melanoma (Supplementary Figure 1c and 2c). As was observed for melanoma, a substantial proportion of causal spatial eQTL-gene interactions in BCC (46%) and SCC (51%) involved non-nearest genes (Supplementary Figure 1d and 2d). Additionally, the causal effect directions were collinear for most (> 90%) of the shared genes in both SCC and BCC (Supplementary Figure 1c and 2c). BCC also showed enrichment for cancer driver genes (*CASP8*, *DDB2*, *FANCA*, *FGFR1OP*, *POU5F1*, *SIRPA*; hypergeometric *P* = 0.028), including specific BCC driver genes (*DDB2*; hypergeometric *P* = 0.039) (Supplementary Table 11), while SCC identified cancer driver genes (*CASP8*, *ARNT*, *DAXX*, *FANCA*, *FCGR2B*, *FUBP1*, *SETDB1*) but did not show enrichment (hypergeometric *P* = 0.19; Supplementary Table 12). Notably, 60% (n = 37) and 87% (n = 109) of the causal genes in BCC and SCC, respectively, are novel and not listed in DisGeNET or the GWAS Catalog (Supplementary Table 13 and 14).

### 3.4 Mendelian randomization detects additional loci beyond those prioritized by GWAS alone

Overall, we identified 144 independent loci implicated in skin cancer risk (Figure 3a, Supplementary Table 1) across 20 chromosomes (Figure 3b). Of these, 33 loci were shared by at least two types of skin cancer, with 15 loci common to all three (Figure 3a and b).

**Figure 3.**
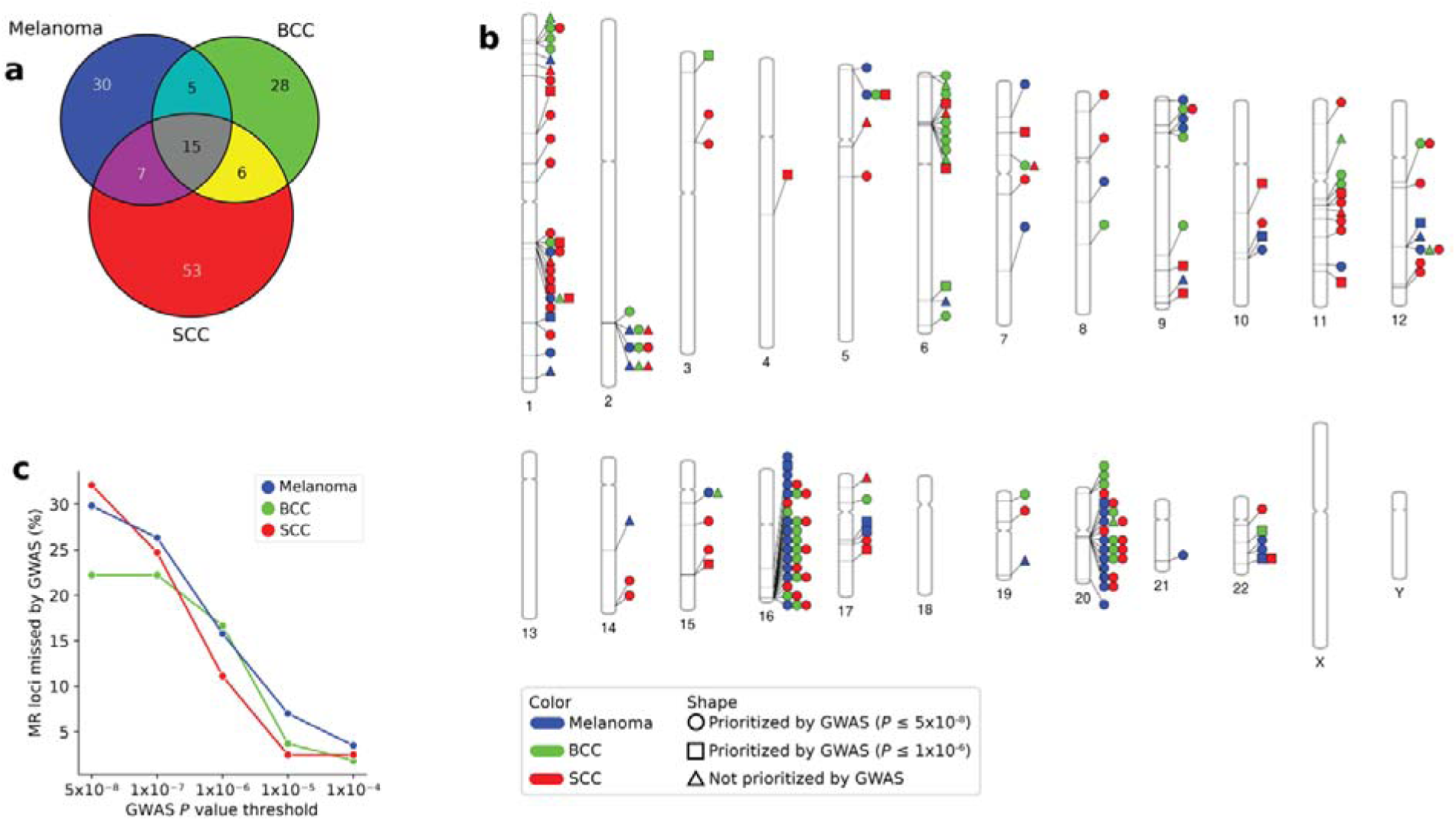
Mendelian randomization (MR) identifies shared and unique causal loci across skin cancer types. (a) Venn diagram showing the unique number of causal loci identified by MR across four tissues in melanoma, basal cell carcinoma (BCC), and squamous cell carcinoma (SCC). (b) Phenogram showing the chromosomal location of MR causal loci identified in each skin cancer type. (c) The percentage of MR causal loci missed by GWAS across increasingly relaxed *P* value thresholds.

For melanoma, MR identified 57 causal loci. Among these, 30% (17/57) were not prioritized by GWAS, as no SNPs within these loci (within LD r^2^ ≥ 0.2) reached genome-wide significance (*P* ≤ 5×10^-8^) in the associated GWAS summary statistics (Figure 3b and c). Additionally, 16% (9/57) were not prioritized even at a GWAS suggestive threshold of *P* ≤ 1×10^-6^.

Similar patterns were observed for BCC and SCC (Figure 3b and c). For BCC, 22% (12/54) of causal loci identified by MR were not prioritized by GWAS at the genome-wide significance level, and 17% (9/54) remain unprioritized at the suggestive threshold. For SCC, 32% (26/81) of causal loci were not prioritized by GWAS at the genome-wide level, and 11% (9/81) were unprioritized at the suggestive threshold.

### 3.5 Mendelian randomization reveal shared and distinct biological bases of skin cancer types

To elucidate the biological consequences of the shared and distinct genetic risks among skin cancer types, we conducted comparative analyses of their causal genes and associated pathways. We identified a total of 206 causal genes, with 49 shared between at least two skin cancer types: 10 genes were exclusively shared between melanoma and BCC, 20 between melanoma and SCC, 5 between BCC and SCC, and 14 were common to all three skin cancer types (Figure 4a). Permutation tests (n = 100,000 simulations) confirmed that the overlap of causal genes between each pair and among all three skin cancer types was significantly greater than expected by chance (Supplementary Figure 3). However, distinct causal gene sets were also characterized for each skin cancer type: 38 genes were specific to melanoma, 33 to BCC, and 86 to SCC (Figure 4a). Pathway enrichment analysis of each gene set revealed unique patterns that putatively highlight the shared and unique biological underpinnings among different skin cancer types (Figure 4b-h, Supplementary Table 15a-g).

**Figure 4.**
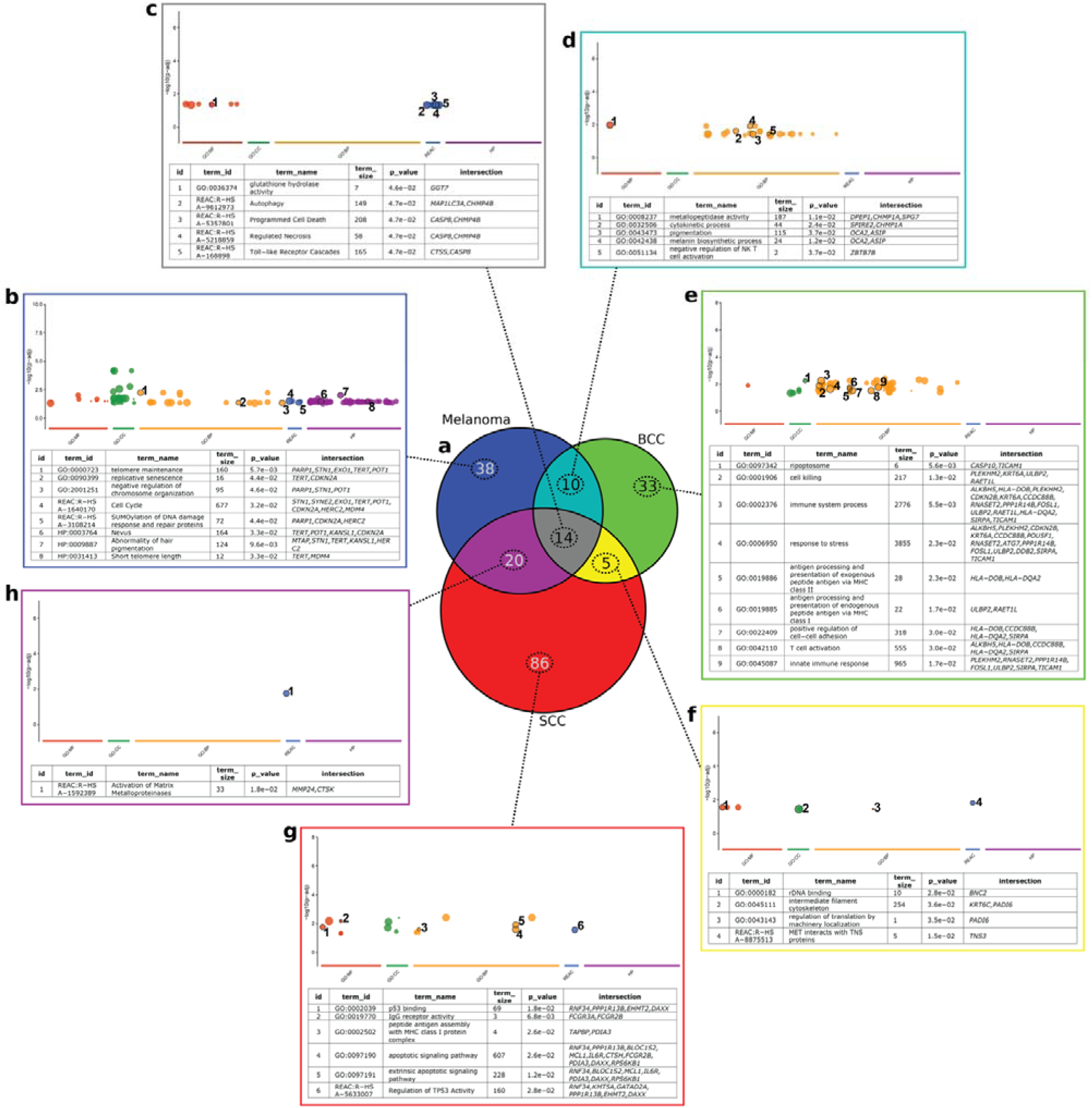
Shared and unique causal genes were identified for each type of skin cancer. (a) Venn diagram showing the number of genes causally associated with melanoma, basal cell carcinoma (BCC), and squamous cell carcinoma (SCC). (b-h) Statistically enriched biological terms (adj. *P* < 0.05) for each gene set are presented in colored rectangles.

Melanoma-specific causal genes were enriched in biological processes including telomere biology and nevus trait (Figure 4b, Supplementary Table 15a). In contrast, genes unique to BCC were predominantly enriched in immune-related processes (Figure 4e, Supplementary Table 15d). Whereas SCC-specific genes were uniquely enriched in p53 biology, in addition to some immune-related processes such as MHC class I protein complex assembly (Figure 4g, Supplementary Table 15f). Additionally, the 10 genes exclusively shared between melanoma and BCC were enriched in pathways such as pigmentation alongside immune processes such as negative regulation of Natural Killer T cell activation (Figure 4d, Supplementary Table 15c).

Finally, genes common among all three skin cancer types were enriched in pathways related to programmed cell death, driven by *CASP8* and *CHMP4B* (Figure 4c, Supplementary Table 15b). Notably, decreased expression of *CASP8*—which is causal across all four tissues of all three skin cancer types—is regulated by four spatial eQTLs across three independent loci (Figure 5). This includes rs62193755 in sun-exposed skin, positioned 268,565 bp downstream from *CASP8*, in the intron of *ALS2CR11* (Figure 5).

**Figure 5.**
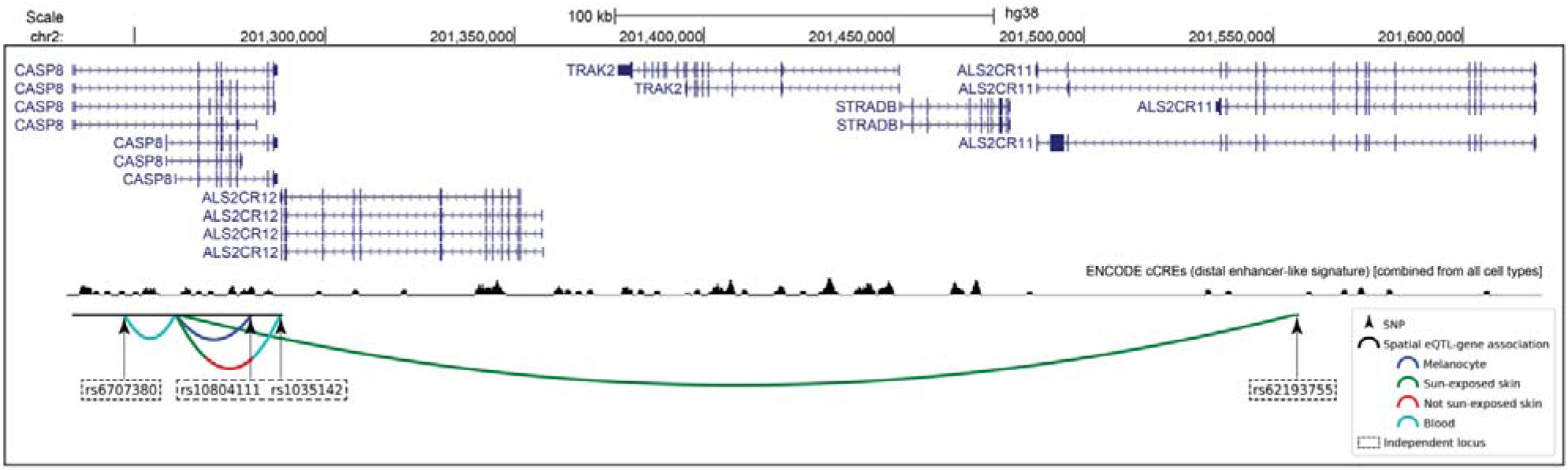
UCSC genome browser tracks spanning chr2:201,231,695-201,620,952. Four spatial eQTLs across three independent loci regulating *CASP8* are depicted. Interactions are indicated for melanocyte (blue), sun-exposed skin (green), not sun-exposed skin (red), and blood (cyan). These specific regulatory interaction patterns are consistent across all skin cancer types.

## 4 Discussion

We conducted two-sample MR analyses to identify causal associations between gene expression and skin cancer outcomes. These analyses integrated the largest GWAS data available on the three major skin cancer types—melanoma, BCC, and SCC—and spatial eQTL data from melanocyte, sun-exposed skin, not sun-exposed skin, and blood.

Across the four tissues, we identified 82, 62, and 125 putative causal genes for melanoma, BCC, and SCC, respectively, many of which were novel associations. We also identified the (spatial eQTL) SNPs regulating these genes and their loci. Notably, many of these MR-identified loci were independent of any GWAS significant SNPs, potentially representing new loci not prioritized by conventional GWAS. This observation mirrors finding by Porcu et al., who reported that 31% of MR loci in their study were missed by GWAS, likely due to power issues^38^. Indeed, as the GWAS sample size increased, fewer MR loci were missed, indicating a saturation effect^38^. Although not directly investigated, it is plausible that similar factors contributed to our observations.

The melanoma and BCC causal gene sets included cancer-specific driver genes (*CDKN2A* and *TERT* for melanoma; *DDB2* for BCC), which showed significant enrichment. Downregulation of all three was associated with increased risk. *DDB2* is essential for the repair of UV-induced DNA damage via the nucleotide excision repair pathway^39^. Germline mutations in *DDB2* are seen in Xeroderma Pigmentosum^40^, a hereditary disease characterized by extreme UV sensitivity and a high predisposition to skin cancers. *CDKN2A*, crucial for cell cycle regulation^41^, is the most commonly somatically inactivated gene in melanoma^42–45^. Germline mutations in *CDKN2A* are seen in Familial Atypical Multiple Mole Melanoma syndrome^46–48^. Thus, the observation that downregulation of *CDKN2A* and *DDB2* was causal for melanoma and BCC risk aligns with their roles as tumor suppressors.

Our analyses revealed that the causal genes identified for melanoma, BCC, and SCC overlap significantly more than is expected by chance, pointing to common underlying pathways. However, we also identified genes specific to each cancer type, highlighting that distinct biological mechanisms may also be at play.

For example, pathway analysis revealed enrichment of telomere biology-related terms among melanoma-specific causal genes. This includes *TERT*, although the observed direction of effect (reduced expression appears to increase risk) deviates from conventional understanding: longer telomeres have been linked to melanoma risk^49^, while frequent somatic mutations in the *TERT* promoter increases its expression^50^. Indeed, the expression changes we observed in several other telomere-related genes, including the downregulation *STN1*^51^, and *MDM4*^52^, and the upregulation of *POT1*^53^, were more consistent with shorter telomeres. In this context, it is worth considering that the loss of telomere protection can lead a state of genome instability, which have been proposed to impose strong selection pressure for stabilization mechanisms, which could make cells more prone to subsequently acquiring the activating *TERT* promoter mutations^54–56^. This raises the possibility that while moderate telomere shortening could be protective through limiting replicative potential, very short telomeres could increase risk by promoting genome instability and conditions that favor the selection of cells that can stabilize it. However, these interpretations remain speculative and further investigations are warranted to clarify the nuanced role of telomere biology in melanoma risk.

It has been suggested that there are at least two pathways for melanoma development (via pigmentation and via nevi) and two for BCC development (via pigmentation and independent of pigmentation)^8^. Our study supports this, as nevus trait is uniquely enriched in melanoma causal genes but not in BCC or SCC. This also aligns with the results of our previous comorbidity analysis, which highlights the divergence between melanoma and other skin cancers due to the unique risk associated with nevus count trait^15^. Additionally, we identified that pigmentation pathways are enriched in the causal genes shared between melanoma and BCC, emphasizing that these skin cancers share a common developmental pathway via pigmentation. This enrichment is driven by *OCA2* (downregulation increase risk) and *ASIP* (upregulation increase risk). *OCA2* plays a role in melanin synthesis^57^, mutations or deletions in *OCA2* are known to cause Oculocutaneous albinism type II^58,59^. *ASIP* encodes the agouti-signaling protein (ASIP), a competitive inhibitor that blocks the UV-stimulated binding of α-MSH to the melanocortin 1 receptor (MC1R)^60^. This inhibition interferes with the UV-induced processes of melanin synthesis and DNA damage repair, which rely on the binding of α-MSH to MC1R^61^. Additionally, ASIP acts as an inverse agonist, decreasing basal MC1R signaling and inhibiting eumelanogenesis^62^. Therefore, the *ASIP* upregulation identified in this study may increase skin cancer risk by reducing MC1R activity and melanin production^63,64^. Of note, ASIP was recently implicated as a potential drug target for both melanoma and BCC^64^.

The interplay between the immune system and tumor, known as immunoediting, is an important factor in cancer development^65^. In particular, BCC-specific causal genes are predominantly enriched in immune-related functions, with enrichment in both MHC class I and class II functions, while SCC-specific genes show enrichment only in MHC class I functions. Additionally, genes shared between BCC and melanoma are enriched for the negative regulation of Natural Killer T cell activation, driven by downregulation of *ZBTB7B*, which is required for the maturation and activation of invariant Natural Killer T (iNKT) cells^66,67^. MHC molecules play a role in eliciting anti-tumor immune responses by presenting endogenous (MHC class I) and exogenous (MHC class II) antigens for immune detection^68^. Additionally, iNKT cells exhibit anti-tumor activity via their cytotoxic and adjuvant activities^69,70^. Together, these findings emphasize the role of immune evasion, a hallmark of cancer^71,72^, in skin cancer development. Notably, genetic susceptibility to BCC appears to be especially influenced by inherited immune system traits.

Regulation of *TP53* activity and p53 binding are specifically enriched for SCC. *TP53* encodes the tumor suppressor protein p53, which is crucial in cancer-related processes such as cell cycle control, DNA repair, apoptosis, and senescence^73^. While p53 is often somatically inactivated in skin cancers^74^, it plays a particularly significant role in the early pathogenesis of SCC, with mutations in *TP53* identified as early events in SCC development^75^. In contrast, early development of BCC appears to be driven by constitutive activation of the sonic hedgehog signaling pathway, primarily due to acquired mutations in the *PTCH* and *SMO* genes^76,77^.

Among the terms enriched across all three skin cancers are those for apoptosis and programmed cell death, driven by *CHMP4B* (upregulated) and *CASP8* (downregulated). High expression of *CHMP4B* is associated with accelerated proliferation and resistance to therapy in hepatocellular carcinoma^78^. *CASP8*, a key initiator of the extrinsic apoptotic pathway, is downregulated in many cancers^79^. Importantly, *CASP8* is causal across all four tissues of all three skin cancer types, being downregulated by four spatial eQTLs across three independent loci. This includes rs62193755 in sun-exposed skin, positioned 268,565 bp downstream from *CASP8*, in the intron of *ALS2CR11*. This suggests a convergence of independent, long-range genomic signals that collectively increase skin cancer risk through *CASP8* downregulation, potentially contributing to apoptosis resistance, a hallmark of cancer^71^.

Several limitations need to be discussed. First, MR assumes that the instrumental variables (spatial eQTLs) are robustly associated with the exposure (gene expression), free from confounders, and influence the outcome (skin cancer) solely through the exposure (*i.e.*, no horizontal pleiotropy)^16^. We selected spatial eQTLs using a stringent threshold (*P* ≤ 1×10[[) as instruments, likely fulfilling the first assumption. The second and third assumptions, however, cannot be empirically proven^80^. Nonetheless, since an individual’s genotype is randomly determined from their parental genotypes at conception, using these randomly allocated genetic variants as instruments should naturally help mitigate confounder effects^16,81^. Furthermore, we performed sensitivity analyses such as the Cochran’s Q test and MR-Egger regression (methods) to remove exposures with potential horizontal pleiotropy^82^.

Second, the GRNs were constructed using Hi-C and eQTL data from different individuals than those in the GWAS datasets. To address this mismatch, we used data from European ancestry populations, which helps mitigate some concerns about the match between variant-exposure and variant-outcome associations. Nonetheless, the focus on European ancestry populations may limit the generalizability of our findings. Thus, further validation is needed to determine whether our findings hold true for other populations or ethnic groups.

Third, the GRNs used in this study only provide a static representation of gene regulatory interactions in specific cell lines at a given time point, potentially overlooking dynamic changes that occur in living systems. Additionally, these GRNs were constructed from a small number of Hi-C and eQTL datasets that may not represent all possible connections. Previous simulation-based analyses have shown that two-sample MR will identify more robust causal relationships as the eQTL sample size increases^38^.

## 5 Conclusion

Despite these limitations, our study provides new insights into the shared and distinct biological mechanisms driving the development of three distinct types of skin cancer. By identifying novel drivers of germline risk, we have provided further avenues for exploration for genetic markers of skin cancer risk and highlighted potential targets for future therapeutic interventions.

## 6 Funding and acknowledgements

JOS was funded by donations from the Dines Family trust. WS was supported by a postdoctoral fellowship from the Vision Research Foundation and a Royal Society of New Zealand Marsden Grant (20-UOA-002). MP was funded by a University of Auckland doctoral scholarship. This work contains data from the Genotype-Tissue Expression (GTEx) Project, which was supported by the Common Fund of the Office of the Director of the National Institutes of Health, and by NCI, NHGRI, NHLBI, NIDA, NIMH, and NINDS.

## 7 Authors’ contributions

MP contributed to conceptualization, performed analyses, data processing, data interpretation, and wrote the manuscript. WS and JOS supervised MP, conceptualized, and co-wrote the manuscript. All authors contributed to the article and approved the submitted version.

## 8 Ethics approval and consent to participate

Ethical approvals and informed consent were obtained by the original data providers for the collection and use of the datasets utilized in this study.

## 9 Consent for publication

Not applicable.

## 10 Data availability

Access to melanoma GWAS summary statistics was approved by the dbGaP Data Access Committee (Project ID: 30073, accession: phs001868.v1.p1). BCC and SCC summary statistics are accessible through GWAS catalog (study accessions: GCST90013410 and GCST90137412, respectively). Access to melanocyte genotype and RNA-seq expression data from 106 individuals was approved by the dbGaP Data Access Committee (Project ID: 30073, accession: phs001500.v1.p1). The CoDeS3D pipeline is available on github (https://github.com/Genome3d/codes3d-v2). GRNs are available on figshare (https://doi.org/10.17608/k6.auckland.27050866.v1). Data analyses and visualizations were performed using Python (version 3.8.12) through Jupyter notebook (version 6.4.6) or using R (version 4.0.4) through RStudio (version 1.4.1106). Additional in-house scripts used for data wrangling are available upon request.

## Supporting information

Supplementary Checklist

## Supplementary Figures

**Supplementary Figure 1.**
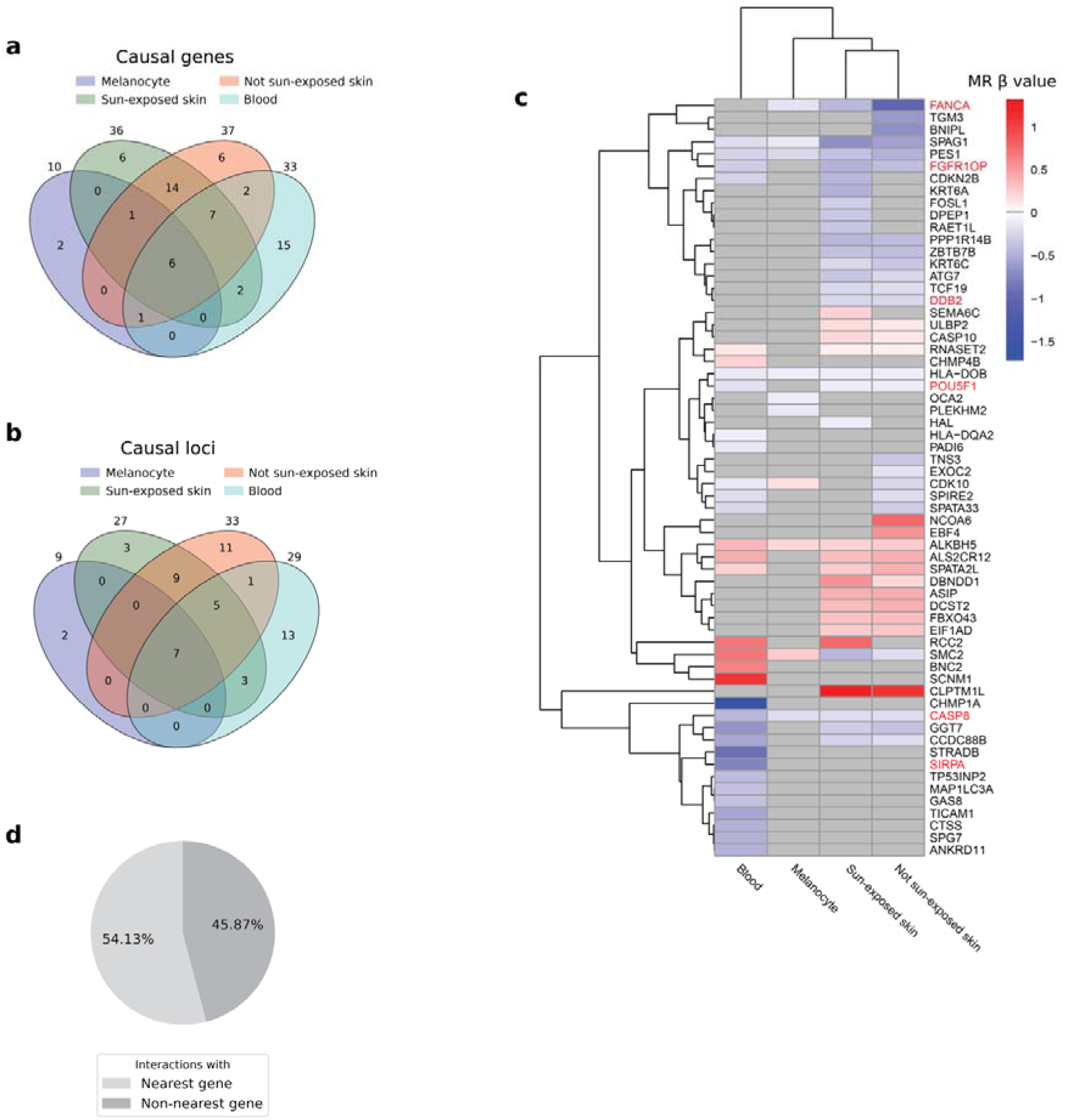
62 genes are causally associated with basal cell carcinoma (BCC) by 2-sample Mendelian randomization (MR). (a) Venn diagram showing the number of BCC causal genes identified in each tissue. (b) Venn diagram showing the number of BCC causal loci in each tissue. (c) Unsupervised hierarchical bi-clustering of the MR β estimates of causal genes across the four tissues. In the context of MR, the β value represents the effect of each unit increase in gene expression on the log(odds) of developing the disease. Therefore, a positive value (red) signifies that increased expression of a gene is associated with elevated BCC risk, whereas a negative value (blue) denotes the opposite effect. Genes highlighted in red are recognized as known cancer drivers in the Cancer Gene Census database. (d) Pie chart showing the proportions of causal SNP-gene interactions that involve nearest genes versus non-nearest genes.

**Supplementary Figure 2.**
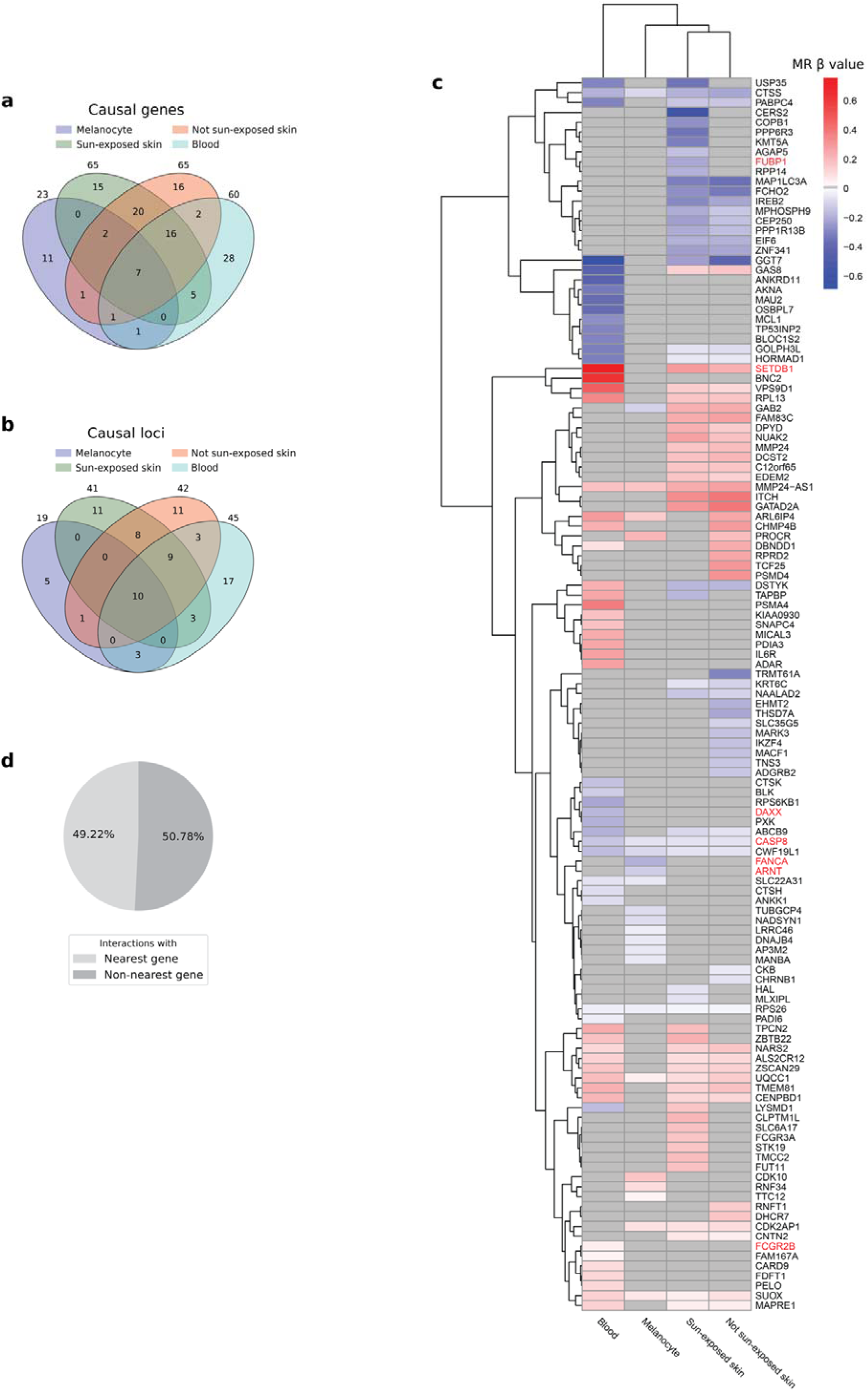
125 genes are causally associated with squamous cell carcinoma (SCC) by 2-sample Mendelian randomization (MR). (a) Venn diagram showing the number of SCC causal genes identified in each tissue. (b) Venn diagram showing the number of SCC causal loci in each tissue. (c) Unsupervised hierarchical bi-clustering of the MR β estimates of causal genes across the four tissues. In the context of MR, the β value represents the effect of each unit increase in gene expression on the log(odds) of developing the disease. Therefore, a positive value (red) signifies that increased expression of a gene is associated with elevated SCC risk, whereas a negative value (blue) denotes the opposite effect. Genes highlighted in red are recognized as known cancer drivers in the Cancer Gene Census database. (d) Pie chart showing the proportions of causal SNP-gene interactions that involve nearest genes versus non-nearest genes.

**Supplementary Figure 3.**
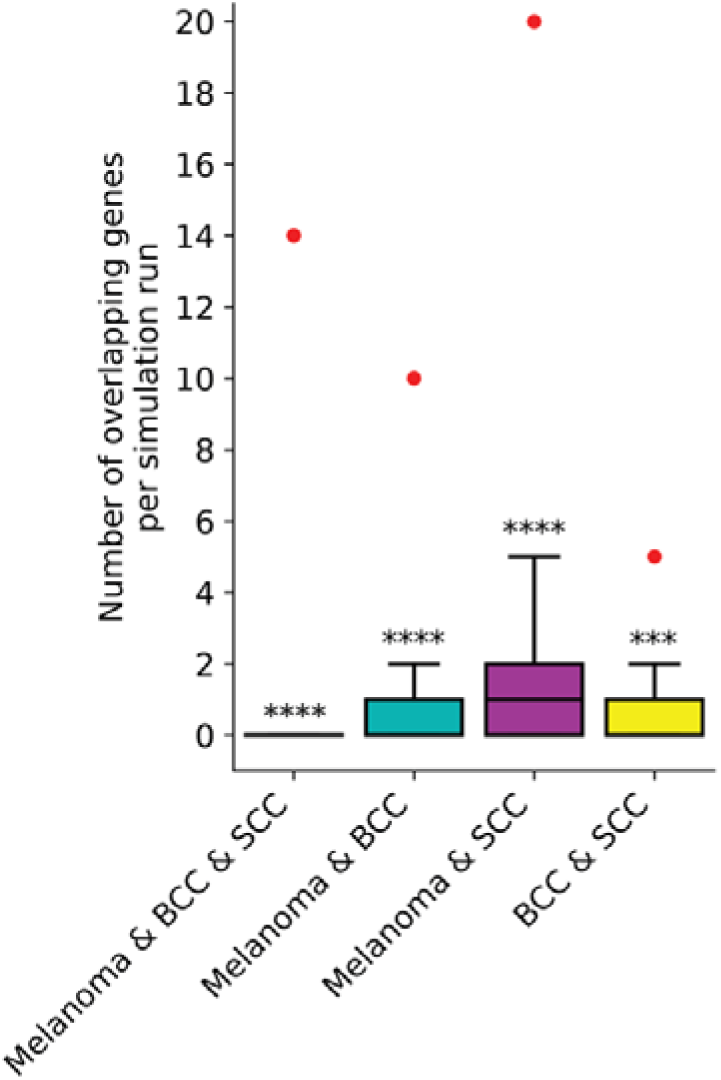
Causal genes of melanoma, basal cell carcinoma (BCC), and squamous cell carcinoma (SCC) overlap significantly more than expected by chance. The observed overlaps from Mendelian randomization analyses (red dots) are significantly above the distribution of overlapping genes from 100,000 bootstrap simulations (boxplots) formed from random gene sets drawn from the 11,833 unique genes in the tissue-specific GRNs. Significance levels are indicated as follows: \**P* ≤ 0.05, \*\**P* ≤ 0.01, \*\*\**P* ≤ 0.001, \*\*\*\**P* ≤ 0.0001.

