## Supplementary Checklist for "Mendelian randomizations with spatial gene networks reveal shared and distinct drivers of risk in major skin cancer types"

**Supplementary Checklist. STROBE-MR checklist of recommended items to address in reports of Mendelian randomization studies**^1,2^**.**

| **Item No.** | **Section** | **Checklist item** | **Page No.** | **Relevant text from manuscript** |
| --- | --- | --- | --- | --- |
| 1 | **TITLE and ABSTRACT** | Indicate Mendelian randomization (MR) as the study’s design in the title and/or the abstract if that is a main purpose of the study | Detailed in title and introduction section | Title: Mendelian randomizations with spatial gene networks reveal shared and distinct drivers of risk in major skin cancer types  Introduction: We then performed two-sample MR, using these spatial eQTLs as instrumental variables, along with GWAS summary statistics for melanoma, BCC and SCC, to infer causal relationships between specific gene expression changes and each type of skin cancer. |
|  | **INTRODUCTION** |  |  |  |
| 2 | **Background** | Explain the scientific background and rationale for the reported study. What is the exposure? Is a potential causal relationship between exposure and outcome plausible? Justify why MR is a helpful method to address the study question | Detailed in introduction section | Genome-wide association studies (GWAS) have identified dozens of genomic loci associated with the risk of developing each type of skin cancer. These GWAS provide a framework for identifying putative risk loci, but due to the nature of these association studies, they are not aimed at identifying causal genes. |
| 3 | **Objectives** | State specific objectives clearly, including pre-specified causal hypotheses (if any). State that MR is a method that, under specific assumptions, intends to estimate causal effects | Detailed in introduction section | We previously investigated how melanoma-associated SNPs influence gene expression by integrating tissue-specific chromatin interaction (Hi-C) and expression quantitative trait loci (eQTL) data. While these findings provided valuable insights, they have yet to establish statistical evidence supporting the causal involvement of these genes in melanoma pathogenesis. Mendelian randomization (MR) is a statistical method that uses genetic variants as instrumental variables to estimate the causal effect of a modifiable exposure on a disease outcome. When appropriately designed, MR is less susceptible to common epidemiological biases and can yield reliable causal inferences.  In this study, we first constructed gene regulatory networks (GRNs) by integrating tissue-specific Hi-C and eQTL data. These GRNs consist of SNPs that physically interact with and regulate the expression of specific target genes, referred to as “spatial eQTLs”. We then performed two-sample MR19, using these spatial eQTLs as instrumental variables, along with GWAS summary statistics for melanoma, BCC and SCC, to infer causal relationships between specific gene expression changes and each type of skin cancer. |
|  | **METHODS** |  |  |  |
| 4 | **Study design and data sources** | Present key elements of the study design early in the article. Consider including a table listing sources of data for all phases of the study. For each data source contributing to the analysis, describe the following: |  |  |
|  | a) | Setting: Describe the study design and the underlying population, if possible. Describe the setting, locations, and relevant dates, including periods of recruitment, exposure, follow-up, and data collection, when available. | Detailed in methods section | Tissue-specific GRNs were generated for four tissues: melanocyte, sun-exposed skin, not sun-exposed skin, and blood using Hi-C data from three skin and four blood cell lines, along with eQTL data from three GTEx and one melanocyte dataset (Supplementary Table 1).  *NOTE : Detailed information and study design of the Hi-C and eQTL data can be found in the cited publications in Supplementary Table 1.*  For melanoma, we used GWAS summary statistics for 36,760 cases and 375,188 controls from Landi et al. as the outcome data. For BCC and SCC, we used GWAS summary statistics from Adolphe et al. (17,416 cases, 375,455 controls) and Seviiri et al. (7,402 cases, 286,892 controls) as the outcome data, respectively.  *NOTE : Detailed information and study design of the GWAS data can be found in the cited GWAS publication.* |
|  | b) | Participants: Give the eligibility criteria, and the sources and methods of selection of participants. Report the sample size, and whether any power or sample size calculations were carried out prior to the main analysis |  | *NOTE : Detailed information and study design of the Hi-C and eQTL data can be found in the cited publications in Supplementary Table 1.*  *NOTE : Detailed information and study design of the GWAS data can be found in the cited GWAS publication.* |
|  | c) | Describe measurement, quality control and selection of genetic variants |  | *NOTE : Detailed information and study design of the Hi-C and eQTL data can be found in the cited publications in Supplementary Table 1.*  *NOTE : Detailed information and study design of the GWAS data can be found in the cited GWAS publication.* |
|  | d) | For each exposure, outcome, and other relevant variables, describe methods of assessment and diagnostic criteria for diseases |  | *NOTE : Detailed information and study design of the Hi-C and eQTL data can be found in the cited publications in Supplementary Table 1.*  *NOTE : Detailed information and study design of the GWAS data can be found in the cited GWAS publication.* |
|  | e) | Provide details of ethics committee approval and participant informed consent, if relevant |  | *NOTE : Detailed information and study design of the Hi-C and eQTL data can be found in the cited publications in Supplementary Table 1.*  *NOTE : Detailed information and study design of the GWAS data can be found in the cited GWAS publication.* |
| 5 | **Assumptions** | Explicitly state the three core IV assumptions for the main analysis (relevance, independence and exclusion restriction) as well assumptions for any additional or sensitivity analysis | Detailed in methods section | The instrumental variables used in MR must satisfy three assumptions: 1) the instrumental variables are associated with the exposure of interest; 2) the instrumental variables are independent of any potential confounders; and 3) the instrumental variables influence the outcome solely through their association with the exposure (i.e., no horizontal pleiotropy). |
| 6 | **Statistical methods: main analysis** | Describe statistical methods and statistics used |  |  |
|  | a) | Describe how quantitative variables were handled in the analyses (i.e., scale, units, model) | - | - |
|  | b) | Describe how genetic variants were handled in the analyses and, if applicable, how their weights were selected | Detailed in methods section | Spatial eQTL-target gene pairs (P ≤ 1x10^-5^) from each GRN were used as the exposure dataset. Palindromic SNPs (i.e., those with alleles that are reverse complements, such as A/T or C/G) were removed to prevent distortion of strand orientation or allele coding during the harmonization process. To ensure the independence of instrumental variables for each exposure, we performed LD clumping within a 10 Mb window and r^2^ cut-off of < 0.001 based on the 1000 Genomes project European population. |
|  | c) | Describe the MR estimator (e.g. two-stage least squares, Wald ratio) and related statistics. Detail the included covariates and, in case of two-sample MR, whether the same covariate set was used for adjustment in the two samples | Detailed in methods section | Once harmonized, MR was performed using the Wald ratio method for exposures with one instrumental variable, inverse variance weighted (IVW) method for exposures with two instrumental variables, and weighted median method for exposures with ≥3 instrumental variables. Target genes with MR *P* value ≤ the Bonferroni-correction threshold of 0.05/(number of unique exposure genes) were considered statistically significant. |
|  | d) | Explain how missing data were addressed | - | - |
|  | e) | If applicable, indicate how multiple testing was addressed | Detailed in methods section | Target genes with MR *P* value ≤ the Bonferroni-correction threshold of 0.05/(number of unique exposure genes) were considered statistically significant. |
| 7 | **Assessment of assumptions** | Describe any methods or prior knowledge used to assess the assumptions or justify their validity | Detailed in methods section | Spatial eQTL-target gene pairs (P ≤ 1x10^-5^) from each GRN were used as the exposure dataset. |
| 8 | **Sensitivity analyses and additional analyses** | Describe any sensitivity analyses or additional analyses performed (e.g. comparison of effect estimates from different approaches, independent replication, bias analytic techniques, validation of instruments, simulations) | Detailed in methods section | We performed several sensitivity analyses for target genes with multiple instrumental variables. First, Cochran’s Q test was performed; instrumental variables with *P* value < 0.05 were considered heterogeneous. For target genes with ≥3 instrumental variables, MR-Egger regression was further performed to detect horizontal pleiotropy. A significant deviation from zero in the intercept (*P* value < 0.05) was taken as evidence of horizontal pleiotropy. Genes failing any of these sensitivity tests were excluded from the list of significant causal genes. |
| 9 | **Software and pre-registration** |  |  |  |
|  | a) | Name statistical software and package(s), including version and settings used | Detailed in methods section | To identify potentially causal genes for each type of skin cancer, we conducted two-sample mendelian randomization (MR) analyses using the TwoSampleMR R package (v0.5.6) |
|  | b) | State whether the study protocol and details were pre-registered (as well as when and where) | - | - |
|  | **RESULTS** |  |  |  |
| 10 | **Descriptive data** |  |  |  |
|  | a) | Report the numbers of individuals at each stage of included studies and reasons for exclusion. Consider use of a flow diagram | Detailed in methods section | *NOTE : Detailed information and study design of the Hi-C and eQTL data can be found in the cited publications in Supplementary Table 1.*  *NOTE : Detailed information and study design of the GWAS data can be found in the cited GWAS publication.* |
|  | b) | Report summary statistics for phenotypic exposure(s), outcome(s), and other relevant variables (e.g. means, SDs, proportions) | Detailed in the methods and data availability sections | Access to melanoma GWAS summary statistics was approved by the dbGaP Data Access Committee (Project ID: 30073, accession: phs001868.v1.p1). BCC and SCC summary statistics are accessible through GWAS catalog (study accessions: GCST90013410 and GCST90137412, respectively). Access to melanocyte genotype and RNA-seq expression data from 106 individuals was approved by the dbGaP Data Access Committee (Project ID: 30073, accession: phs001500.v1.p1).  *NOTE: Access to all other publicly available data are detailed on Supplementary Table 1.* |
|  | c) | If the data sources include meta-analyses of previous studies, provide the assessments of heterogeneity across these studies | - | *NOTE : Detailed information and study design of the Hi-C and eQTL data can be found in the cited publications in Supplementary Table 1.*  *NOTE : Detailed information and study design of the GWAS data can be found in the cited GWAS publication.* |
|  | d) | For two-sample MR:  i.  Provide justification of the similarity of the genetic variant-exposure associations between the exposure and outcome samples  ii.  Provide information on the number of individuals who overlap between the exposure and outcome studies |  | *NOTE : Detailed information and study design of the Hi-C and eQTL data can be found in the cited publications in Supplementary Table 1.*  *NOTE : Detailed information and study design of the GWAS data can be found in the cited GWAS publication.* |
| 11 | **Main results** |  |  |  |
|  | a) | Report the associations between genetic variant and exposure, and between genetic variant and outcome, preferably on an interpretable scale |  | Supplementary File 1 |
|  | b) | Report MR estimates of the relationship between exposure and outcome, and the measures of uncertainty from the MR analysis, on an interpretable scale, such as odds ratio or relative risk per SD difference |  | Supplementary Table 4, 8, and 10 |
|  | c) | If relevant, consider translating estimates of relative risk into absolute risk for a meaningful time period | - | - |
|  | d) | Consider plots to visualize results (e.g. forest plot, scatterplot of associations between genetic variants and outcome versus between genetic variants and exposure) | - | - |
| 12 | **Assessment of assumptions** |  |  |  |
|  | a) | Report the assessment of the validity of the assumptions | Detailed in methods section and supplementary tables | We selected spatial eQTLs using a stringent threshold (*P* ≤ 1x10⁻⁵) as instruments, likely fulfilling the first assumption. The second and third assumptions, however, cannot be empirically proven. Nonetheless, since an individual’s genotype is randomly determined from their parental genotypes at conception, using these randomly allocated genetic variants as instruments should naturally help mitigate confounder effects. Furthermore, we performed sensitivity analyses such as the Cochran’s Q test and MR-Egger regression (methods) to remove exposures with potential horizontal pleiotropy.  *NOTE : The result of sensitivity analyses are reported in Supplementary Table 3, 7, and 9* |
|  | b) | Report any additional statistics (e.g., assessments of heterogeneity across genetic variants, such as *I^2^*, Q statistic or E-value) |  | *Supplementary Table 3, 7, and 9* |
| 13 | **Sensitivity analyses and additional analyses** |  |  |  |
|  | a) | Report any sensitivity analyses to assess the robustness of the main results to violations of the assumptions |  | *Supplementary Table 3, 7, and 9* |
|  | b) | Report results from other sensitivity analyses or additional analyses |  | *Supplementary Table 3, 7, and 9* |
|  | c) | Report any assessment of direction of causal relationship (e.g., bidirectional MR) | - | - |
|  | d) | When relevant, report and compare with estimates from non-MR analyses | - | - |
|  | e) | Consider additional plots to visualize results (e.g., leave-one-out analyses) | - | - |
|  | **DISCUSSION** |  |  |  |
| 14 | **Key results** | Summarize key results with reference to study objectives | Detailed in discussion section | Despite these limitations, our study provides new insights into the shared and distinct biological mechanisms driving the development of three distinct types of skin cancer. By identifying novel drivers of germline risk, we have provided further avenues for exploration for genetic markers of skin cancer risk and highlighted potential targets for future therapeutic interventions. |
| 15 | **Limitations** | Discuss limitations of the study, taking into account the validity of the IV assumptions, other sources of potential bias, and imprecision. Discuss both direction and magnitude of any potential bias and any efforts to address them | Detailed in discussion section | Several limitations need to be discussed. First, MR assumes that the instrumental variables (spatial eQTLs) are robustly associated with the exposure (gene expression), free from confounders, and influence the outcome (skin cancer) solely through the exposure (i.e., no horizontal pleiotropy). We selected spatial eQTLs using a stringent threshold (*P* ≤ 1x10⁻⁵) as instruments, likely fulfilling the first assumption. The second and third assumptions, however, cannot be empirically proven. Nonetheless, since an individual’s genotype is randomly determined from their parental genotypes at conception, using these randomly allocated genetic variants as instruments should naturally help mitigate confounder effects. Furthermore, we performed sensitivity analyses such as the Cochran’s Q test and MR-Egger regression (methods) to remove exposures with potential horizontal pleiotropy.  Second, the GRNs were constructed using Hi-C and eQTL data from different individuals than those in the GWAS datasets. To address this mismatch, we used data from European ancestry populations, which helps mitigate some concerns about the match between variant-exposure and variant-outcome associations. Nonetheless, the focus on European ancestry populations may limit the generalizability of our findings. Thus, further validation is needed to determine whether our findings hold true for other populations or ethnic groups.  Third, the GRNs used in this study only provide a static representation of gene regulatory interactions in specific cell lines at a given time point, potentially overlooking dynamic changes that occur in living systems. Additionally, these GRNs were constructed from a small number of Hi-C and eQTL datasets that may not represent all possible connections. Previous simulation-based analyses have shown that two-sample MR will identify more robust causal relationships as the eQTL sample size increases. |
| 16 | **Interpretation** |  |  |  |
|  | a) | Meaning: Give a cautious overall interpretation of results in the context of their limitations and in comparison with other studies | Detailed in discussion section | Several limitations need to be discussed. First, MR assumes that the instrumental variables (spatial eQTLs) are robustly associated with the exposure (gene expression), free from confounders, and influence the outcome (skin cancer) solely through the exposure (i.e., no horizontal pleiotropy). We selected spatial eQTLs using a stringent threshold (*P* ≤ 1x10⁻⁵) as instruments, likely fulfilling the first assumption. The second and third assumptions, however, cannot be empirically proven. Nonetheless, since an individual’s genotype is randomly determined from their parental genotypes at conception, using these randomly allocated genetic variants as instruments should naturally help mitigate confounder effects. Furthermore, we performed sensitivity analyses such as the Cochran’s Q test and MR-Egger regression (methods) to remove exposures with potential horizontal pleiotropy.  Second, the GRNs were constructed using Hi-C and eQTL data from different individuals than those in the GWAS datasets. To address this mismatch, we used data from European ancestry populations, which helps mitigate some concerns about the match between variant-exposure and variant-outcome associations. Nonetheless, the focus on European ancestry populations may limit the generalizability of our findings. Thus, further validation is needed to determine whether our findings hold true for other populations or ethnic groups.  Third, the GRNs used in this study only provide a static representation of gene regulatory interactions in specific cell lines at a given time point, potentially overlooking dynamic changes that occur in living systems. Additionally, these GRNs were constructed from a small number of Hi-C and eQTL datasets that may not represent all possible connections. Previous simulation-based analyses have shown that two-sample MR will identify more robust causal relationships as the eQTL sample size increases. |
|  | b) | Mechanism: Discuss underlying biological mechanisms that could drive a potential causal relationship between the investigated exposure and the outcome, and whether the gene-environment equivalence assumption is reasonable. Use causal language carefully, clarifying that IV estimates may provide causal effects only under certain assumptions | Detailed in discussion section | Discussed in detail throughout the discussion section. |
|  | c) | Clinical relevance: Discuss whether the results have clinical or public policy relevance, and to what extent they inform effect sizes of possible interventions | Detailed in discussion section | By identifying novel drivers of germline risk, we have provided further avenues for exploration for genetic markers of skin cancer risk and highlighted potential targets for future therapeutic interventions. |
| 17 | **Generalizability** | Discuss the generalizability of the study results (a) to other populations, (b) across other exposure periods/timings, and (c) across other levels of exposure | Detailed in discussion section | Second, the GRNs were constructed using Hi-C and eQTL data from different individuals than those in the GWAS datasets. To address this mismatch, we used data from European ancestry populations, which helps mitigate some concerns about the match between variant-exposure and variant-outcome associations. Nonetheless, the focus on European ancestry populations may limit the generalizability of our findings. Thus, further validation is needed to determine whether our findings hold true for other populations or ethnic groups.  Third, the GRNs used in this study only provide a static representation of gene regulatory interactions in specific cell lines at a given time point, potentially overlooking dynamic changes that occur in living systems. |
|  | **OTHER INFORMATION** |  |  |  |
| 18 | **Funding** | Describe sources of funding and the role of funders in the present study and, if applicable, sources of funding for the databases and original study or studies on which the present study is based | Detailed in funding and acknowledgements section | JOS was funded by donations from the Dines Family trust. WS was supported by a postdoctoral fellowship from the Vision Research Foundation and a Royal Society of New Zealand Marsden Grant (20-UOA-002). MP was funded by a University of Auckland doctoral scholarship. |
| 19 | **Data and data sharing** | Provide the data used to perform all analyses or report where and how the data can be accessed, and reference these sources in the article. Provide the statistical code needed to reproduce the results in the article, or report whether the code is publicly accessible and if so, where | Detailed in data availability section | Access to melanoma GWAS summary statistics was approved by the dbGaP Data Access Committee (Project ID: 30073, accession: phs001868.v1.p1). BCC and SCC summary statistics are accessible through GWAS catalog (study accessions: GCST90013410 and GCST90137412, respectively). Access to melanocyte genotype and RNA-seq expression data from 106 individuals was approved by the dbGaP Data Access Committee (Project ID: 30073, accession: phs001500.v1.p1). The CoDeS3D pipeline is available on github (https://github.com/Genome3d/codes3d-v2). GRNs are available on figshare (https://doi.org/10.17608/k6.auckland.27050866.v1). Data analyses and visualizations were performed using Python (version 3.8.12) through Jupyter notebook (version 6.4.6) or using R (version 4.0.4) through RStudio (version 1.4.1106). Additional in-house scripts used for data wrangling are available upon request.  *NOTE: Access to all other publicly available data are detailed on Supplementary Table 1.* |
| 20 | **Conflicts of Interest** | All authors should declare all potential conflicts of interest | Detailed in competing interests section | All authors have seen and approved the final manuscript. They do not have any competing interests to declare. |

This checklist is copyrighted by the Equator Network under the Creative Commons Attribution 3.0 Unported (CC BY 3.0) license.
